# Incomplete transcripts dominate the *Mycobacterium tuberculosis* transcriptome

**DOI:** 10.1101/2023.03.10.532058

**Authors:** Xiangwu Ju, Shuqi Li, Ruby Froom, Ling Wang, Mirjana Lilic, Elizabeth A. Campbell, Jeremy M. Rock, Shixin Liu

## Abstract

*Mycobacterium tuberculosis* (Mtb) is a bacterial pathogen that causes tuberculosis, an infectious disease that inflicts major health and economic costs around the world^1^. Mtb encounters a diversity of environments during its lifecycle, and responds to these changing environments by reprogramming its transcriptional output^2^. However, the transcriptomic features of Mtb remain poorly characterized. In this work, we comprehensively profile the Mtb transcriptome using the SEnd-seq method that simultaneously captures the 5’ and 3’ ends of RNA^3^. Surprisingly, we find that the RNA coverage for most of the Mtb transcription units display a gradual drop-off within a 200-500 nucleotide window downstream of the transcription start site, yielding a massive number of incomplete transcripts with heterogeneous 3’ ends. We further show that the accumulation of these short RNAs is mainly due to the intrinsically low processivity of the Mtb transcription machinery rather than trans-acting factors such as Rho. Finally, we demonstrate that transcription-translation coupling plays a critical role in generating full-length protein-coding transcripts in Mtb. In sum, our results depict a mycobacterial transcriptome that is dominated by incomplete RNA products, suggesting a distinctive set of transcriptional regulatory mechanisms that could be exploited for new therapeutics.

## Main

Tuberculosis (TB) is the leading cause of death among infectious diseases^4^. The etiological agent of TB, *Mycobacterium tuberculosis* (Mtb), has an exceptional ability to evade host defense and drug treatment^1^. Mtb achieves this feat by enacting coordinated transcriptional control of its gene expression in response to changing environments^2^. Thus, a detailed characterization of the Mtb transcriptome in different genetic backgrounds and under diverse growth conditions is key to understanding the pathogenesis and persistence of TB. However, previous studies suggest that the operation of the Mtb transcription machinery differs significantly from its well-studied *Escherichia coli* (Eco) counterpart^5,6^ and many aspects of Mtb transcription remain poorly understood.

Recently, we developed an RNA profiling method, SEnd-seq, to simultaneously determine the 5’- and 3’-end sequences of the same RNA molecules in a bacterial transcriptome^3^. This method provides a greater resolution of transcript boundaries than standard RNA-seq and generated new insights into the mechanism of transcription in model organisms such as Eco^3^. In this work, we applied SEnd-seq in combination with chemical and genetic manipulation and in vitro biochemistry to characterize the properties of the Mtb transcriptome and transcription machinery.

### Profiling the Mtb transcriptome by SEnd-seq

Compared to Eco, Mtb has a much slower growth rate and a more complex cell wall. To adapt SEnd-seq to Mtb, we modified the protocols for cell disruption, RNA isolation and library preparation (Extended Data Fig. 1 and Methods). Our method yielded, for the first time, correlated positions of the 5’ and 3’ termini of individual Mtb transcripts (Fig. 1a, b, Extended Data Fig. 2a, b). The resultant Mtb transcriptome revealed several notable features. First, we identified 8,873 transcriptional start sites (TSSs) in Mtb, twice as many as found in Eco, even though these two bacteria have a similar genome size (Fig. 1c, Supplementary Table 1). We also applied SEnd-seq to the non-pathogenic model mycobacterium, *Mycobacterium smegmatis* (Msm)^7^, and found that this fast-growing mycobacterium utilizes fewer TSSs than Mtb despite having a larger genome (Fig. 1c). This higher-than-expected abundance of TSS in Mtb can be attributed to elevated levels of intragenic transcription initiation as well as antisense transcription (Extended Data Fig. 2c-e). SEnd-seq also identified 747 leaderless TSSs that lack a 5’ untranslated region (Extended Data Fig. 2f, g), which add to the known leaderless TSS list in the Mtb transcriptome^8^. Second, we identified very few transcription termination sites (TTSs)—defined here as sites where a sudden drop in the RNA 3’-end level was observed—199 in Mtb compared to 1,285 in Eco and 263 in Msm (Fig. 1d, Extended Data Fig. 3, Supplementary Table 2). The lack of intrinsic terminators in the Mtb genome is consistent with bioinformatic predictions^9^. Third, we detected pervasive antisense RNAs (asRNAs) in the Mtb transcriptome (Extended Data Fig. 4, Supplementary Table 3). More than half of Mtb genes feature at least one asRNA (defined by a unique TSS) in their coding region (Fig. 1e). These findings demonstrate the utility of SEnd-seq to comprehensively profile mycobacterial transcriptomes.

**Figure 1.**
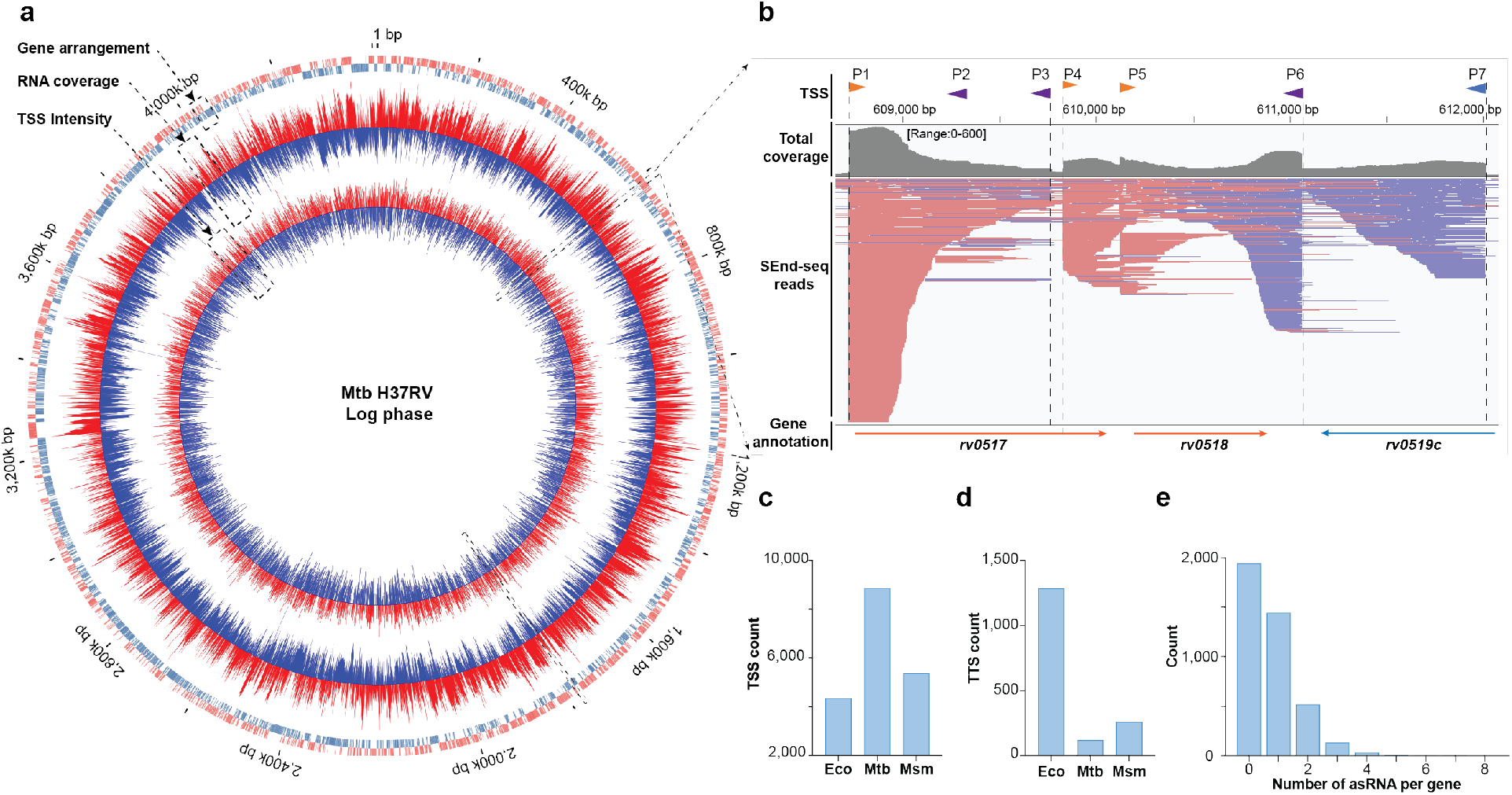
Mtb transcriptome profiling by SEnd-seq. **a**, Circos plot showing the transcriptomic profile of log-phase Mtb cells. Outer circle: gene annotation; middle circle: RNA coverage; inner circle: TSS intensity. Red and blue represent positive and negative strands, respectively. **b**, SEnd-seq data track for an example Mtb genomic region showing total RNA coverage (summed from signals on both strands), aligned SEnd-seq reads (red lines: positive-strand RNA; blue lines: negative-strand RNA), and TSSs (sense: orange arrows; antisense: purple arrows). **c**, Bar graph showing the number of TSSs detected by SEnd-seq for Eco, Mtb and Msm. **d**, Bar graph showing the number of TTSs detected by SEnd-seq for Eco, Mtb and Msm. **e**, Histogram showing the number of asRNAs associated within each Mtb coding gene.

### The Mtb transcriptome is dominated by short incomplete transcripts

Upon examining the Mtb SEnd-seq datasets, we found a strikingly prevalent pattern that the sense RNA level gradually declines between 200 and 500 nucleotides (nt) downstream of its TSS, resulting in an ensemble of transcripts with uniform 5’ ends and heterogeneous 3’ ends (Fig. 1b, 2a). This pattern contrasts with the Eco transcriptome, where the RNA coverage downstream of TSS is typically maintained at a stable level (Fig. 2b). To characterize this drop-off pattern, we analyzed 1,930 protein-coding transcription units (TUs), each utilizing a major TSS that controls one or multiple co-directional genes (Extended Data Fig. 5). To quantify the magnitude of reduction in RNA coverage, we defined a numerical “processivity factor” (PF) as the ratio of the average RNA coverage of a downstream zone over that of an upstream zone for a given TU (Fig. 2c). Consistent with visual inspection, the PF values are predominantly distributed below 1.0 and vary considerably among TUs (Fig. 2d, Extended Data Fig. 6a, b, Supplementary Table 4). TUs with a high PF (>0.5) are enriched in genes that encode components of the Mtb cell envelope for reasons that remain to be determined (Extended Data Fig. 6c). The RNA coverage drop-off was observed for both log- and stationary-phase Mtb cells, with the latter exhibiting a steeper decline (Extended Data Fig. 6d). We then analyzed the Mtb asRNAs and found that they also displayed a drop-off pattern with a mean PF of 0.15 (as compared to 0.47 for coding TUs) (Fig. 2e, Extended Data Fig. 7).

**Figure 2.**
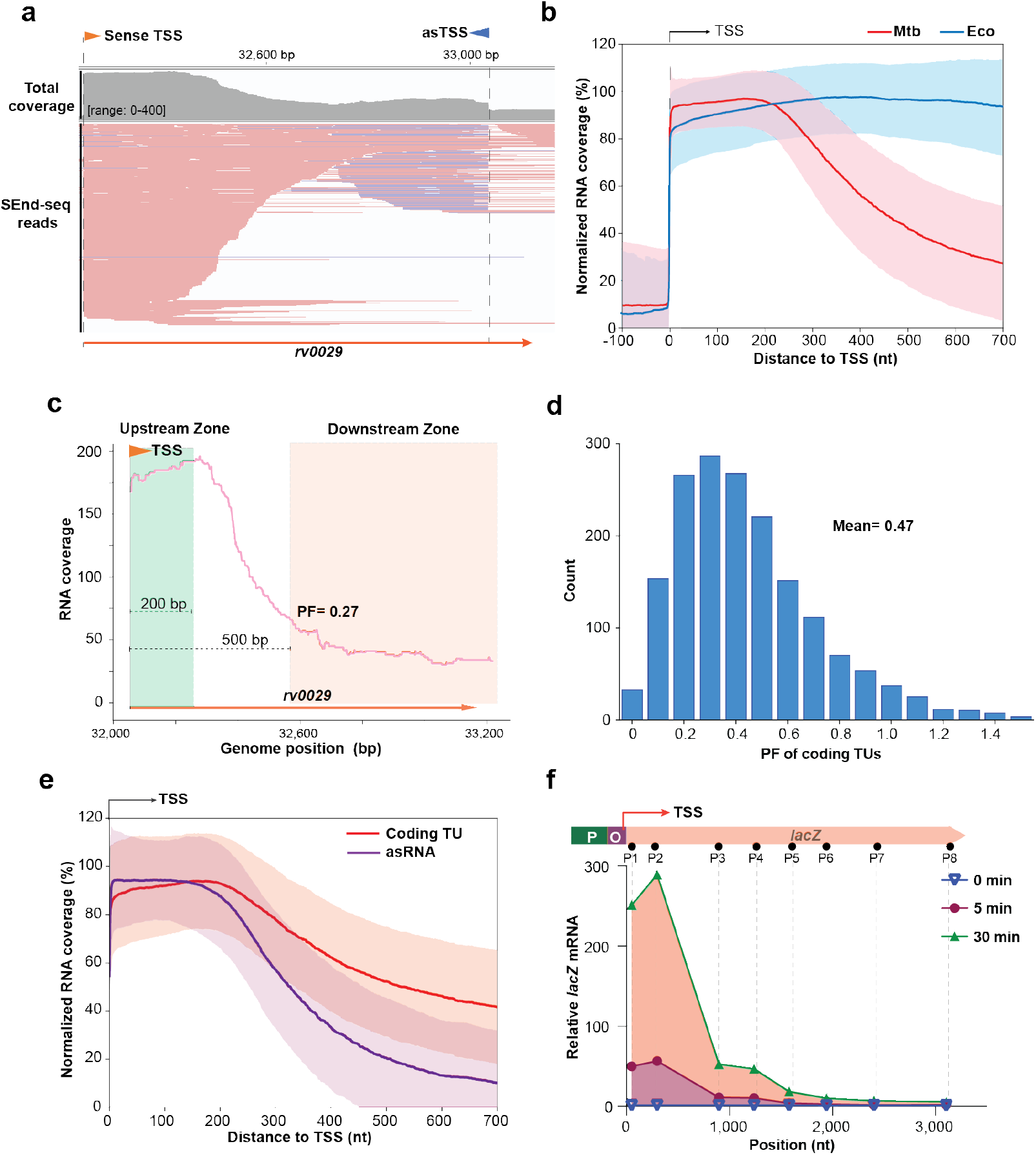
Incomplete RNAs dominate the Mtb transcriptome. **a**, SEnd-seq data track for an example Mtb transcription unit showing predominantly incomplete RNAs accumulating downstream of the TSS (red lines: sense transcripts; blue lines: antisense transcripts). **b**, Summed RNA coverage aligned at TSSs for log-phase Mtb and Eco cells. TSSs without another nearby TSS (±700 nt) were selected for analysis (1,432 sites for Mtb, 932 for Eco). The RNA coverage for each TSS was normalized to the maximum value within a 300-nt region downstream of the TSS. Colored lines (red: Mtb; blue: Eco) represent median values and shaded regions represent standard deviations. **c**, SEnd-seq signals for an example Mtb coding TU demonstrating the definition of upstream zone and downstream zone for PF calculation. **d**, Histogram showing the distribution of PF values for coding TUs (longer than 700 bp) from log-phase Mtb cells. **e**, Summed RNA coverage aligned at TSSs for highly expressed coding TUs (red) and asRNAs (blue) (1,494 coding TUs, 1,553 asRNAs). Colored lines represent median values and shaded regions represent standard deviations. **f**, mRNA length profile for *lacZ* transcription in Mtb at different time points post-induction.

Next, we employed an anhydrotetracycline (ATc)-inducible expression system to monitor real-time RNA synthesis of the heterologous *lacZ* gene in Mtb^10^. We found that upon ATc induction, the RNA abundance proximal to the *lacZ* TSS increased steadily with time, whereas the RNA abundance in the downstream region exhibited minimal increase (Extended Data Fig. 8). The RNA abundance profile obtained 30 minutes after induction shows a drop-off pattern that mirrors the SEnd-seq results (compare Fig. 2f and Extended Data Fig. 6d). Therefore, the short RNAs in the Mtb transcriptome are more likely generated during nascent RNA synthesis than during post-transcriptional RNA processing.

### Mtb Rho is not responsible for short RNA accumulation

Next, we examined whether Rho, the universal transcription termination factor in bacteria^11^, contributes to the generation of incomplete transcripts in Mtb. To this end, we used CRISPR interference (CRISPRi)^12^ to knockdown the expression of *rho* in Mtb, which was confirmed by SEnd-seq (RNA level) and western blot (protein level) (Extended Data Fig. 9a, b). Consistent with the known essentiality of Rho in Mtb^13^, *rho* knockdown significantly impaired cell growth and changed the expression level of many essential genes (Extended Data Fig. 9c, d). However, SEnd-seq analysis showed that *rho* knockdown did not significantly change the PF for coding TUs (Extended Data Fig. 9e), suggesting that the high fraction of incomplete transcripts is not caused by Rho-mediated premature termination. Instead, we found that a general effect of *rho* knockdown was a genome-wide increase in asRNA abundance (without an obvious change in transcript length) (Extended Data Fig. 9f, g).

### Mtb transcription machinery inherently produces short transcripts

We then asked if incomplete transcript production was an inherent property of the Mtb transcription machinery. To this end, we analyzed genome-wide Mtb RNA polymerase (RNAP) occupancy using published RpoB ChIP-seq data^6^ and found that the average RNAP intensity aligned at TSSs displayed a drop-off within a few hundred basepairs downstream of TSS, matching the RNA coverage profile from SEnd-seq (Fig. 3a, b). Based on this observation, we hypothesized that the incomplete RNAs may originate from an intrinsically low processivity of the Mtb transcription complex, which tends to stall at promoter-proximal regions. To test this hypothesis, we performed in vitro transcription experiments using purified Mtb RNAP/σ^A^ holoenzymes and a plasmid DNA template containing a promoter, a 2,435-bp transcribed region, and an intrinsic terminator. We then used qPCR to analyze the RNA abundance at various locations within the gene body as a function of time. Consistent with the in vivo SEnd-seq results, in vitro reactions produced mostly transcripts shorter than 500 nt and few full-length transcripts (Fig. 3c). Similar results were obtained with two other DNA templates (Extended Data Fig. 10a, b). In contrast, the Eco RNAP/σ^70^ holoenzyme synthesized predominantly full-length transcripts in vitro (Fig. 3d). Next, we constructed a template that contained a pre-formed DNA bubble and an RNA primer, which enabled the RNAP to circumvent the need for a σ factor to synthesize RNA. Interestingly, we found that the Mtb RNAP core enzyme alone synthesized full-length transcripts on this template (Extended Data Fig. 10c), but the addition of σ^A^ restored the accumulation of short RNAs (Extended Data Fig. 10d). Together, these results suggest that the prevalent incomplete RNAs observed in Mtb cells can be attributed to the intrinsic processivity of the Mtb transcription machinery, and that the σ factor plays a determining role in such low processivity.

**Figure 3.**
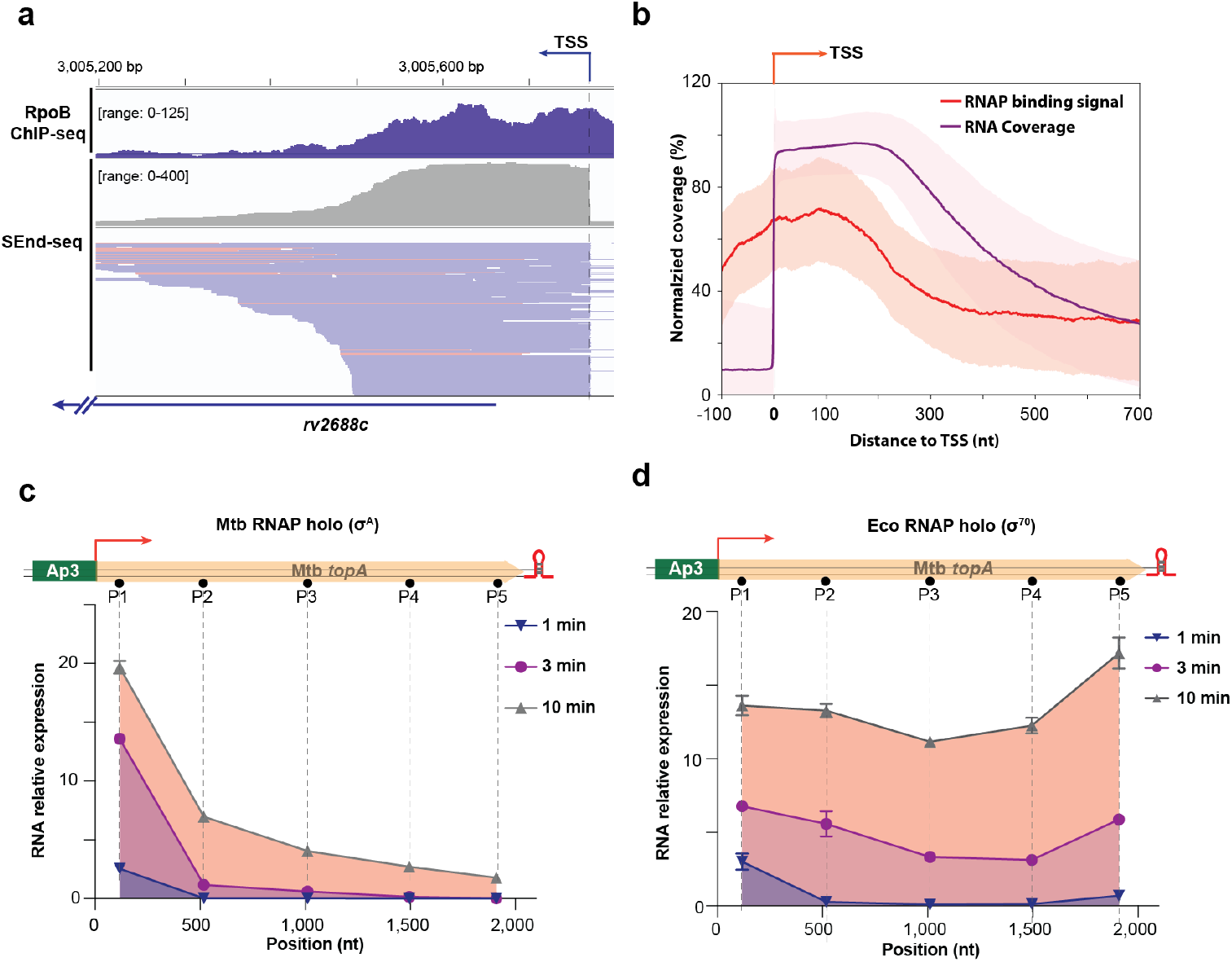
Mtb RNAP/σ^A^ holoenzyme produces mostly short transcripts. **a**, Mtb RNAP ChIP-seq (using antibody against RpoB) and SEnd-seq data tracks for an example Mtb genomic region. **b**, Summed intensities of RNAP occupancy (RpoB ChIP-seq signal; red line) and RNA coverage (SEnd-seq signal; blue line) aligned at Mtb TSSs. **c**, Length profile of RNA products from in vitro transcription assays using Mtb RNAP/σ^A^ holoenzyme and a plasmid DNA template measured by qPCR at different subregions of the template. **d**, Length profile of RNA products from in vitro transcription assays using Eco RNAP/σ^70^ holoenzyme and the same DNA template as in **c**.

### Inhibition of translation exacerbates short RNA accumulation in Mtb

It is known that active translation can facilitate transcription elongation in bacteria^14^. However, the occurrence and functional consequences of transcription-translation coupling in Mtb have remained unclear. Given the poor intrinsic processivity of the Mtb transcription machinery documented here, we speculated that translation may play a key role in synthesizing full-length Mtb transcripts. The steeper drop-off in SEnd-seq coverage for asRNAs than for protein-coding RNAs (Fig. 2e) hinted that ribosome translation may promote Mtb’s transcriptional processivity. We thus analyzed published Mtb ribosome profiling data^15^ and indeed found a strong correlation between the summed ribosome binding signal over a coding TU and the RNA abundance in its downstream zone (Fig. 4a, b). We then treated Mtb cells with the translation inhibitor linezolid^16^ and performed SEnd-seq transcriptomic profiling for these cells. We found that linezolid treatment caused a dramatic reduction in RNA coverage in the downstream zone for coding TUs (Fig. 4c and Extended Data Fig. 11a-c) and shifted the PF distribution towards lower values (Fig. 4d). A similar trend was observed in Mtb cells treated with another ribosome-targeting antibiotic, clarithromycin^17^ (Extended Data Fig. 11a-c). On the other hand, streptomycin, an antibiotic that induces translational miscoding but does not inhibit translation initiation^18^, did not significantly affect the PF of coding TUs (Extended Data Fig. 11d-f). Notably, linezolid treatment had little effect on the transcriptional processivity of asRNAs (Extended Data Fig. 12a), which displayed low ribosome binding signals as expected (Extended Data Fig. 12b, c). Moreover, we used the inducible *lacZ* system to probe the acute response of transcriptional processivity to translation perturbation. We treated Mtb cells with linezolid immediately before ATc induction and found that the RNA abundance in the downstream zone (>300 nt after TSS) was reduced compared to the DMSO-treated control group (Fig. 4e). Finally, we introduced nonsense mutations to the *lacZ* gene body and observed a substantial reduction in RNA abundance downstream of the ectopic stop codons (Fig. 4f). Together, these results demonstrate that active translation by ribosomes aids Mtb transcription elongation and promotes the synthesis of messenger RNAs that encode full-length proteins.

**Figure 4.**
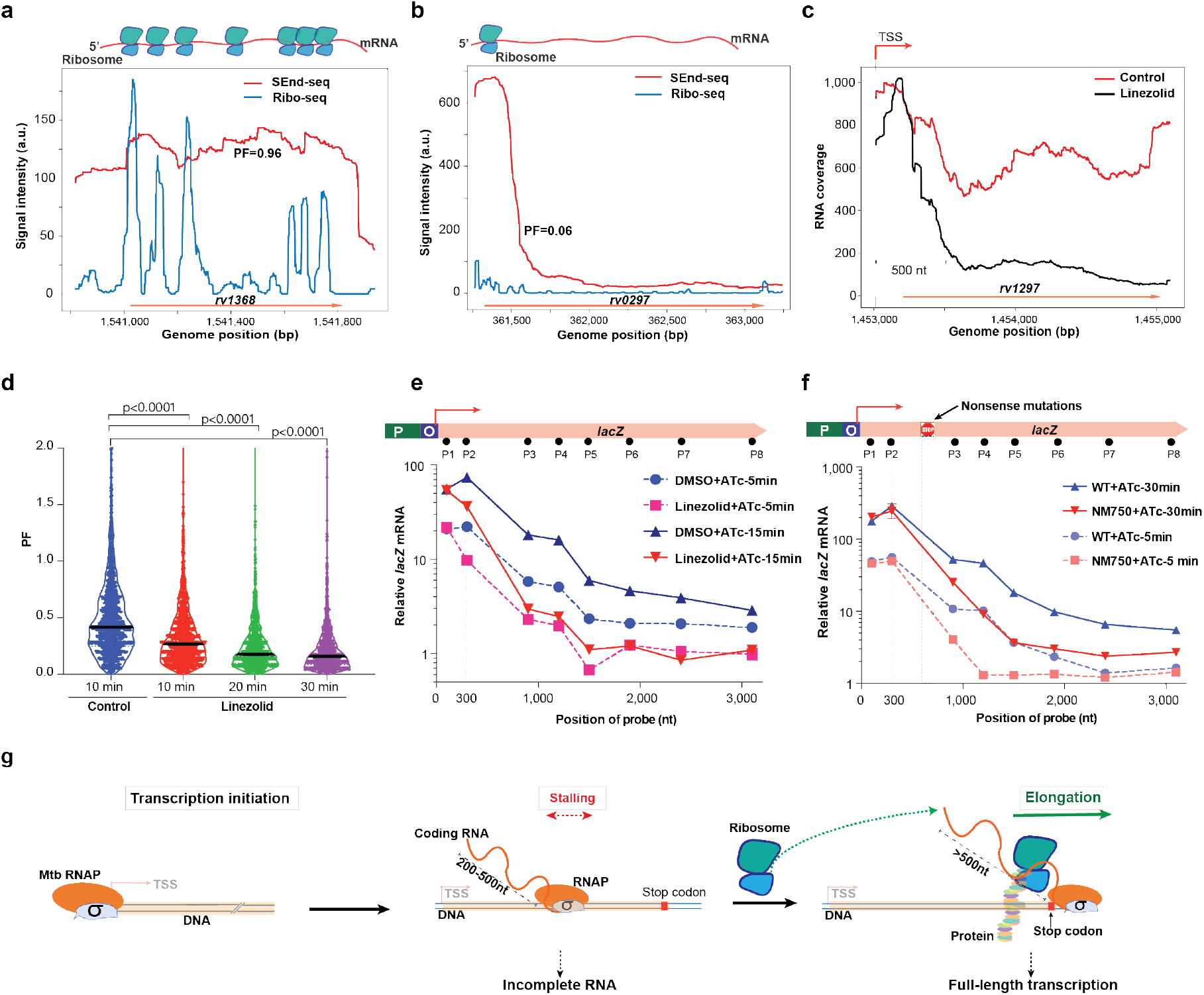
Active translation is important for transcription elongation in Mtb. **a**, SEnd-seq signals (red) for a high-PF Mtb coding TU and the corresponding Ribo-seq signals (blue) in the same region. **b**, SEnd-seq and Ribo-seq signals for a low-PF Mtb coding TU. **c**, SEnd-seq signals in an example Mtb TU showing the difference in RNA coverage between the linezolid-treated condition (black) and the DMSO-treated control condition (red). **d**, Violin plot showing the distribution of PFs for RNA samples collected from Mtb cells treated with linezolid for various periods of time. **e**, Length profile of mRNA products from chromosome-integrated *lacZ* transcription in log-phase Mtb cells induced by ATc and treated with DMSO or linezolid. The RNA level was normalized to the uninduced and DMSO-treated condition. **f**, Length profile of mRNA products from a wildtype (WT) *lacZ* template or a mutant template (NM750) containing two nonsense mutations 750 nt downstream of the TSS. The RNA level was normalized to the WT uninduced condition. The error bars are smaller than the symbols. **g**, Working model of Mtb transcription showing σ-dependent TSS-proximal stalling and translation-stimulated elongation.

## Discussion

Characterization of Mtb transcription and its regulation is critical for understanding how Mtb adapts to diverse environments. Using an RNA profiling method capable of simultaneously capturing 5’ and 3’ ends of the same transcripts, our work reveals that the Mtb transcriptome is dominated by incomplete RNA products accumulating 200-500 nt downstream of TSS. We show that this prevalent pattern is largely due to the inherently low processivity of the Mtb transcription machinery when the RNAP is associated with the housekeeping σ^A^ factor (Fig. 4g). As such, besides its well-known function in promoter recognition and transcription initiation, σ^A^ impedes Mtb RNAP elongation reminiscent of the promoter-proximal pausing during eukaryotic Pol II transcription^19^. Notably, σ-dependent pausing has also been documented in Eco^20^, and Eco RNAP can retain σ^70^ throughout the transcription cycle in some contexts^21^. How σ factors differentially affect the structure and function of Mtb transcription machinery warrants further investigation.

Transcription-translation coupling is a hallmark of bacterial gene expression^22^. However, its functional relevance varies among species^10,23^ and has remained uncharacterized in mycobacteria. Our data suggest that, for a major fraction of Mtb coding genes, ribosome activity is crucial for the transcription elongation complex to synthesize full-length mRNA (Fig. 4g). This may provide an evolutionary advantage to ensure that cells do not waste energy on synthesizing unused full-length transcripts. Interestingly, such polarity effect in Mtb differs from that in Eco in that it does not depend on the activity of Rho^24^, but instead is implemented by the transcription machinery itself. For TUs with a low PF, other gene regulatory mechanisms such as riboswitches are likely at work to suppress ribosome binding to the Mtb mRNA^25,26^.

In sum, our study depicts a mycobacterial transcriptome that drastically differs from the Eco paradigm, revealing Mtb-specific regulatory mechanisms that could be exploited for TB therapy. For example, given the tight coupling between Mtb RNAP and ribosome, simultaneously targeting both machines with antibiotics may yield synergistic effects. Genes with a high PF (i.e., more full-length products) are likely more susceptible to such dual-drug intervention, thereby conferring specificity. Moreover, our work serves as a benchmark for dissecting transcriptional reprogramming of Mtb under stress conditions inside the host or upon drug treatment, which may inform new therapeutic avenues.

## Acknowledgements

We thank Seth Darst, Robert Landick, Anna Marie Pyle, and Dirk Schnappinger for sharing reagents, Michael DeJesus and Rui Gong for technical assistance, and Gabriella Chua for a critical reading of the manuscript. This work was supported by the National Institutes of Health (NIH) grant No. 5R01GM114450 (to E.A.C.), the Rita Allen Foundation (to J.M.R.), an NIH New Innovator Award 1DP2AI144850-01 (to J.M.R.), the Robertson Foundation (to S.Liu), the Alfred P. Sloan Foundation (to S.Liu), and an NIH New Innovator Award 1DP2HG010510-01 (to S.Liu).

## Author Contributions

S.Liu, J.M.R. and X.J. conceived the project. X.J. performed the experiments and data analysis. S.Li and J.M.R. provided resources for mycobacterial sample collection. R.F., L.W., M.L. and E.A.C. provided resources for in vitro transcription experiments. X.J. and S.Liu wrote the manuscript with inputs from all authors.

## Methods

### Bacterial strains and growth conditions

*Mycobacterium tuberculosis* (Mtb) H37Rv was grown at 37°C in a minimal medium [Difco Middlebrook 7H9 broth (BD, 271310) supplemented with 0.5% (v/v) glycerol, 0.05% (v/v) tyloxapol, 0.2 g l^-1^ casamino acids, 0.024 g l^-1^ pantothenic acid and 10% (v/v) OADC (Oleic acid, Albumin, Dextrose and Catalase) (BD, 212351)]. The double auxotrophic Mtb mc^2^6206 strain (H37Rv *ΔpanCD ΔleuCD*)^27^ was grown in the minimal medium with an additional 50 mg l^-1^ L-leucine (Sigma, L8000) and 24 mg l^-1^ pantothenic acid (Sigma, P5155). *Mycobacterium smegmatis* (Msm) MC^2^155 was grown in the Middlebrook 7H9 medium supplemented with 0.2% (v/v) glycerol, 0.05% (v/v) Tween-80, and 10% (v/v) albumin-dextrose-catalase. When applicable, 30 μg ml^-1^ linezolid (Sigma, PZ0014), 40 μg ml^-1^ clarithromycin (Sigma, C9742), 300 μg ml^-1^ streptomycin (Sigma, S9137), 100 ng ml^-1^ anhydrotetracycline (Sigma, 37919), or 20 μg ml^-1^ kanamycin (Goldbio, K-120) was added. Mtb and Msm liquid cultures were grown at 37°C in Nalgene sterile square PETG media bottles with constant agitation. Solid cultures of Mtb were grown on 7H10 agar supplemented as described above except for tyloxapol.

### CRISPRi

Plasmid pIRL58 (Addgene, 166886) bearing the *Streptococcus thermophilus* CRISPR-dCas9 system (dCas9_Sth1_)^12^ was used to modulate the RNA expression level of target genes in Mtb mc^2^6206 cells. Oligos for sgRNAs were synthesized by Integrated DNA Technologies and subsequently cloned into pIRL58. After verification by Sanger sequencing, pIRL58 and pIRL19 (Addgene, 163634), which supplied L5 integrase function on a separate suicide vector, were co-transformed into Mtb cells by electroporation using GenePulser (BioRad) at 2,500 V, 700 ohm and 25 μF. Single colonies were picked from the solid culture plates with 20 μg ml^-1^ kanamycin selection after 14-21 days of culture. Target gene knockdown was induced by adding 100 ng ml^-1^ ATc. The sgRNA and primer sequences are listed in Supplementary Table 5.

### SEnd-seq experiments

#### RNA isolation

Bacterial cells were quenched by adding 1 × vol of GTC buffer [600 g l^-1^ guanidium thiocyanate, 5 g l^-1^ N-laurylsarcosine, 7.1 g l^-1^ sodium citrate, and 0.7% (v/v) β-mercaptoethanol] to the culture medium immediately before harvest and placed at room temperature for 15 min. Cell pellets were collected by centrifugation (4,000 g for 10 min at 4 °C), then thoroughly resuspended in 100 μl of TE buffer (10 mM Tris-HCl pH 8.0 and 1 mM EDTA). After adding 1 ml of TRIzol Reagent (Invitrogen, 15596) and 300 mg of glass beads (Sigma, G1145), the cells were immediately lysed by bead beating with the Precellys Evolution homogenizer (Bertin Technologies, 02520-300-RD000) at 10,000 rpm for 4× 45-s cycles with intervals of 60 s and chilled with dry ice. After removal of the beads by spinning samples at 12,000 rpm for 5 min at 4°C, the liquid phase was transferred to a new tube. 200 μl of chloroform was added, and the sample was gently inverted several times until reaching homogeneity. The sample was then incubated for 15 min at room temperature before spinning at 12,000 g for 10 min at 4°C. The upper phase (~600 μl) was gently collected and mixed at a 1:1 ratio with 100% isopropanol. The sample was incubated for 1 h at −20°C and then centrifuged at 14,000 rpm for 15 min at 4°C. The pellet was washed twice with 1 ml of 75% (v/v) ethanol, air dried for 5 min, and dissolved in nuclease-free water. RNA integrity was assessed with 1% (m/v) agarose gel and Agilent 2100 Bioanalyzer System (Agilent Technologies, 5067-4626). For antibiotic treatment conditions, Mtb mc^2^6206 cells were exponentially grown to an OD_600_ of ~0.8 followed by treatment with indicated antibiotic at specific concentrations. At each time point following the treatment, 4 ml of cell culture medium was withdrawn and mixed with 4 ml of GTC buffer quickly. The cells were then harvested, and the RNA was isolated as described above.

#### Library preparation for total RNA SEnd-seq

5 μg of total RNA was mixed with pooled spike-in RNA used in our previous study^3^ at a mass ratio of 300:1 in a total volume of 12 μl. The RNA sample was incubated with a 5’ adaptor ligation mix [1 μl of 100 μM 5’ adaptor (Supplementary Table 5), 0.5 μl of 50 mM ATP, 2 μl of dimethyl sulfoxide (DMSO), 5 μl of 50% PEG8000, 1 μl of RNase Inhibitor (New England BioLabs, M0314), and 1 μl of High Concentration T4 RNA Ligase 1 (New England BioLabs, M0437)] at 23°C for 5 h. The sample was then diluted with nuclease-free water and cleaned twice with 1.5× vol of Agencourt RNAClean XP beads (Beckman Coulter, A63987). Immediately following this step, the eluted RNA was ligated to the 3’ adaptor (Supplementary Table 5) using the same procedure as for 5’ adaptor ligation. After incubation at 23°C for 5 h, the reaction was diluted to 40 μl with water and purified twice with 1.5× vol of Agencourt RNAClean XP beads to remove excess adaptors. The sample was subsequently eluted with 0.1× TE buffer and subjected to ribosomal RNA removal with RiboMinus Transcriptome Isolation Kit (ThermoFisher, K155004) following the manufacturer’s instructions. After recovery by ethanol precipitation, the RNA was reverse transcribed to cDNA with *Eubacterium rectale* maturase (recombinantly purified from Eco, a gift from A. M. Pyle, Yale University)^28^ and 5’-phosphorylated and biotinylated reverse transcription primer (Supplementary Table 5). After purification, the cDNA was circularized with TS2126 RNA Ligase^29^ (a gift from K. Ryan, City College of New York). Double-stranded DNA was synthesized with DNA Polymerase I (New England BioLabs, M0209S) and subsequently fragmented by dsDNA Fragmentase (New England BioLabs, M0348S) at 37°C for 15 min. The reaction was stopped by adding 5 μl of 0.5 M EDTA and incubated at 65°C for 15 min in the presence of 50 mM DTT. Next, the DNA was diluted to 40 μl with TE buffer and purified with 1.5× vol of AMPure beads (Beckman Coulter, A63882). The eluted DNA was used for sequencing library preparation with NEBNext Ultra II DNA Library Prep Kit (New England BioLabs, E7645). Biotinylated DNA fragments were enriched by Dynabeads M-280 Streptavidin (ThermoFisher, 11205D) and further amplified for 12 cycles by PCR.

#### Library preparation for primary RNA SEnd-seq

5 μg of total RNA was used for primary transcript enrichment with our previously published method^3^. Briefly, the 5’-PPP RNA in the total RNA was specifically capped with 3’-Desthiobiotin-GTP (New England BioLabs, N0761) by the Vaccinia Capping System (New England BioLabs, M2080S). The RNA was subjected to 3’ adaptor ligation using the same procedure as described above and subsequently enriched with Hydrophilic Streptavidin Magnetic Beads (New England BioLabs, S1421). After washing thoroughly, the RNA was eluted and reverse transcribed to cDNA as described above. The remaining steps were the same as those for library preparation for total RNA SEnd-seq, except that the DNA library was amplified for 15 cycles.

#### Illumina sequencing

Following PCR amplification, each amplicon was cleaned by 1× vol of AMPure XP beads twice and quantified with a Qubit 2.0 fluorometer (Invitrogen). The amplicon size and purity were further evaluated on an Agilent 2200 Tape Station (Agilent Technologies, 5067-5576). Equal amounts of amplicon were then multiplexed and sequenced with 2× 150 cycles on an Illumina MiSeq, NextSeq500, or NovaSeq6000 platform (Rockefeller University Genomics Resource Center).

### SEnd-seq data analysis

#### Data processing

Raw data were processed as previously described^3^. Briefly, pair-end sequences were merged to single-end ones by FLASh software. Correlated 5’ and 3’ end sequences were extracted by identifying the sequence of the adaptor ligated to the 3’ end. Full-length sequences were inferred by mapping the end reads to the corresponding reference genome (NC000913.3 for Eco, NC008596.1 for Msm, and NC018143.2 for Mtb) using Bowtie 2. Reads with an insert length greater than 10,000 nt were discarded. Aligned full-length reads were visualized using the Integrative Genome Viewer.

#### RNA coverage

Each full-length read was first mapped to the genome in a specific direction. Directional RNA coverage was quantified by summing the number of aligned reads at each mapped nucleotide position. RNA coverage for a gene or TU was quantified by summing the coverage of all nucleotide positions spanned by the region of interest. When comparing RNA coverage between samples, the data were normalized by the total non-ribosomal RNA amount from the entire transcriptome. For the samples treated with linezolid, the abundance of spike-in RNA was used for normalization.

#### TSS identification

The TSS identification criteria were the same as we used before for Eco TSS identification^3^. Candidate TSS positions within five nucleotides in the same orientation were grouped together, and the position with the largest amount of read increase was used as the representative TSS position.

#### TTS identification

Potential TTSs were identified from the total RNA SEnd-seq data at nucleotide positions with more than 10 reads and with a reduction of more than 40% in read coverage from its upstream to its downstream.

#### TU annotation

The genome was at first segmented into preliminary TUs which contained annotated genes of the same direction. A preliminary unit was further segmented into multiple units if it contained any internal TSS with a strong activity (>2-fold increase in RNA coverage between downstream and upstream). As such, each TU contains a major TSS (TU start site) and possibly additional minor TSSs (<2-fold increase in RNA coverage). The end site of a TU was set to 10 bp before the start of a following co-directional TU, or the middle position between opposite genes that belong to two convergent TUs.

#### Antisense transcript annotation

asRNAs were called if there existed a strong antisense TSS (asRNA start site) in a given TU and an opposite direction TSS within the non-annotated 400 bp downstream of the coding region. The end site of an asRNA was set to the position where the RNA coverage dropped to 25% of the peak value.

#### PF analysis

Each coding TU was divided into an upstream zone (from 0 to 200 bp downstream of TSS) and a downstream zone (from 500 bp downstream of TSS to the end of TU). If there was another qualified TSS located within the downstream zone, the region downstream of that TSS was excluded from analysis. The ratio of average RNA intensity of the downstream zone over that of the upstream zone was calculated as the processivity factor (PF) for the corresponding TU. For asRNAs, the upstream zone and downstream zone were defined as 0-200 bp downstream of TSS and 500-700 bp downstream of TSS, respectively.

### Ribo-seq data analysis

Mtb ribosome profiling data were downloaded from the EMBL-EBI database (E-MTAB-8835). After read extraction and quality filter, the reads were mapped to the Mtb genome using Bowtie 2. The directional ribosome binding signal was extracted and plotted using a custom script.

### ChIP-seq data analysis

Mtb RpoB ChIP-seq data were downloaded from NCBI GEO (GSE40862). After read extraction and quality filter, the reads were mapped to the Mtb genome using Bowtie 2. The RNAP binding signal was extracted and plotted using a custom script.

### Immunoblot

Mtb cells were lysed with TRIzol Reagent as in *RNA isolation* described above, and protein samples were extracted following a Trizol-based protein extraction protocol provided by the manufacturer. Immunoblotting was performed as described previously^30^. Antibodies against Mtb Rho (a gift from D. Schnappinger, Weill Cornell Medicine) and against Eco RpoB (BioLegend, W0023) were used.

### qPCR

1-10 μg of total RNA was treated with 0.5 μl of TURBO DNase (Life Technologies, AM2238) at 37°C for 30 min to remove the genomic DNA. The RNA was diluted to 100 μl with RNase-free water and then cleaned three times with 100 μl of H_2_O-saturated phenol:chloroform:isoamyl alcohol (25:24:1, v/v/v). After ethanol precipitation, 1 μg of RNA was reverse transcribed to cDNA with the High-capacity cDNA Reverse Transcription Kit (ThermoFisher, 4368814) following the manufacturer’s instructions. qPCR was conducted using synthesized primers and the SYBR green master mix (ThermoFisher, 4309155) on a QuantStudio 6 Flex Real-Time PCR System (Thermo Fisher Scientific). The relative RNA abundance was presented as the signal ratio between the target transcript and the reference 16S rRNA from the same sample using the formula: 2^Ct(16S)-Ct(target)^, where Ct denotes the cycle threshold.

### Inducible *lacZ* transcription in Mtb

Plasmid pIRL58 was modified by removing the sgRNA expression cassette and replacing the dCas9_Sth1_ gene body with the Eco *lacZ* coding region, allowing the synthesis of *lacZ* RNA under the control of ATc-inducible promoter *P_tet_*. The modified plasmid was co-transformed into Mtb mc^2^6206 cells with pIRL19 as described above. Cells from a single colony of Mtb *P_tet_-lacZ* after selection were exponentially grown to an OD_600_ of ~0.8 followed by the addition of 100 ng ml^-1^ ATc to induce *lacZ* transcription. After induction, 4 ml of cell culture was withdrawn at indicated time intervals and mixed with 4 ml of GTC buffer in a new tube as sample *t* (St). One extra sample taken immediately before ATc addition was referred to as S0. After RNA isolation and TURBO DNase treatment as described above, 1 μg of total RNA was used to synthesize the cDNA for qPCR. The relative *lacZ* mRNA abundance at each time point is defined as 2^Ct(S0)-Ct(St)^, where Ct denotes the cycle threshold.

### In vitro transcription

DNA templates were amplified by PCR from Mtb genomic DNA with primer sets listed in Supplementary Table 5. An AP3 promoter sequence was inserted into one end of the template and an intrinsic terminator (derived from TsynB in pIRL58) was placed at the other end. The DNA fragment was then incorporated into the pUC19 plasmid. The plasmid templates were prepared from Eco DH5α cells and subsequently treated with 2 μl RNase A (ThermoFisher, EN0531) for 30 min and 2 μl Proteinase K (New England BioLabs, P8107S) for 1 h. The plasmid templates were further cleaned three times with phenol:chloroform:isoamyl alcohol (25:24:1, v/v/v) and recovered by ethanol precipitation.

To prepare the bubble template, the DNA fragment containing the AP3 promoter at its 5’ end and the intrinsic terminator at its 3’ end was amplified from the plasmid DNA described above by PCR. The product was cleaned with QIAQuick PCR purification kit (Qiagen, 28104) and phenol:chloroform:isoamyl alcohol (25:24:1, v/v/v). The bubble template was constructed by ligating a DNA adaptor (NEBNext adaptor for Illumina) to both ends of the DNA fragment using NEBNext Ultra II DNA Library Prep Kit. After XbaI digestion (cut site immediately after the terminator), the DNA template was purified using AMPure XP beads.

Purified Mtb RNAP/σ^A^ holoenzyme was prepared as described previously^31^. The in vitro transcription mixture contained 2 μl of 10× transcription buffer (100 mM Tris-HCl pH 7.9, 0.5 M KCl, 100 mM MgCl2, 10 mM DTT, 50 μg/mL BSA), 1 μl of RNase Inhibitor, 0.5 pmol of DNA template and 2 pmol of Mtb RNAP holoenzyme (or core RNAP only) in a 20-μl volume. The mixture was incubated at 37°C for 15 min before the addition of rNTPs (100 μM each). At indicated time points, the reaction was quenched by adding 20 mM EDTA (final concentration) and 2 μl of Proteinase K and incubating for 30 min. The reaction was then diluted to 100 μl with RNase-free H2O and cleaned three times with phenol:chloroform:isoamyl alcohol (25:24:1, v/v/v). After ethanol precipitation and resuspension with 30 μl of RNase-free H2O, 0.5 μl of DNase I (New England BioLabs, M0303S), 3.5 μl of DNase buffer and 1 μl of RNase inhibitor were added. After incubation at 37°C for 30 min, the RNA product was cleaned three times with phenol:chloroform:isoamyl alcohol (25:24:1, v/v/v) and recovered by ethanol precipitation. Half of the RNA was evaluated by 5% urea polyacrylamide gel electrophoresis, stained by SYBR Gold Nucleic Acid Gel Stain (ThermoFisher Scientific, S11494), scanned by Axygen Gel Documentation System (Corning, GD1000), and quantified by ImageJ. The remaining half was converted to cDNA with High-capacity cDNA Reverse Transcription Kit and evaluated by qPCR as described above. The RNA abundance was normalized to diluted plasmid DNA template (0.033 ng ml^-1^).

### Data availability

### Code availability

### Statistical analysis

Data are shown as mean ± s.d. unless noted otherwise. *P* values were determined by two-sided unpaired Student’s t-tests using GraphPad Prism 9. The difference between two groups was considered statistically significant when the *P* value is less than 0.05 (\**P* < 0.05; \*\**P* < 0.01; \*\*\**P* < 0.001; \*\*\*\**P* < 0.0001; ns, not significant).

