## Supplementary Figures for "Incomplete transcripts dominate the *Mycobacterium tuberculosis* transcriptome"

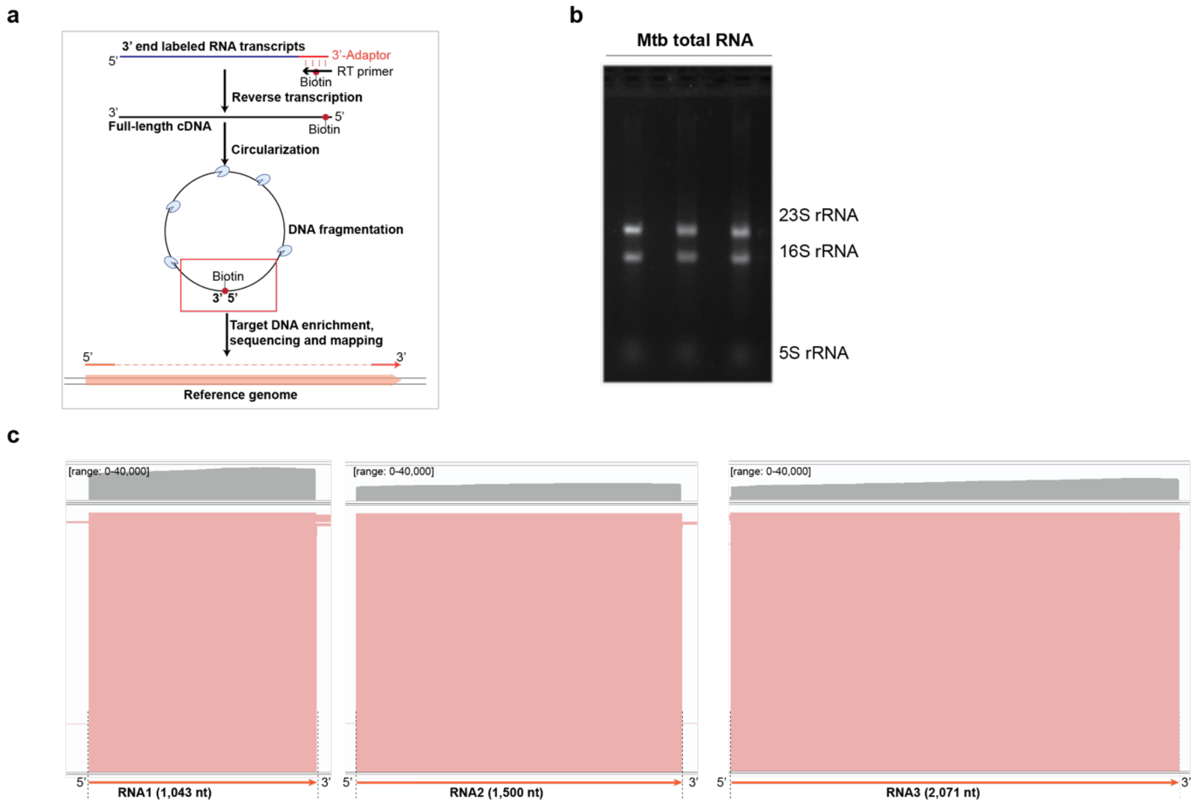


**Extended Data Figure 1. Workflow and quality control of Mtb SEnd-seq. a**, Workflow of Mtb SEnd-seq. See Methods for details. **b**, Gel showing the isolated total RNA from Mtb cells. **c**, SEnd-seq data tracks of three spike-in RNAs with different lengths, which were pooled with total cellular RNA before the preparation of sequencing libraries.

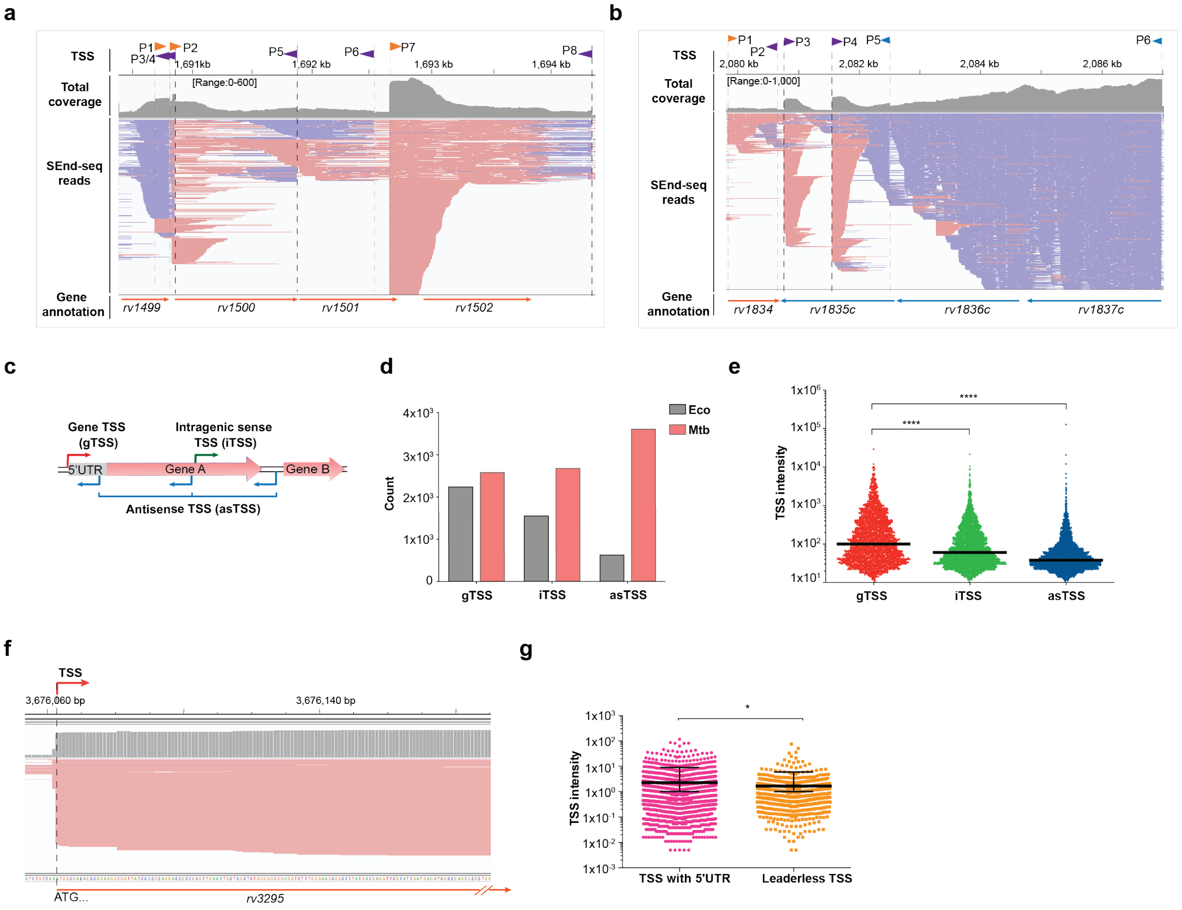


**Extended Data Figure 2. Characterization of Mtb TSSs detected by SEnd-seq. a-b**, SEnd-seq data tracks for two example genomic regions from log-phase Mtb cells. **c**, Schematic showing different categories of TSSs based on their location and orientation. **d**, Number of TSSs in different categories annotated in **c** for Eco and Mtb. **e**, Distribution of Mtb TSS intensities for different categories annotated in **c**. The black bars indicate mean values. **f**, SEnd-seq data track showing an example of leaderless TSS in Mtb. **g**, Distribution of Mtb TSS intensities for TSSs with or without a 5' untranslated region. The black bars indicate mean values and standard deviations.

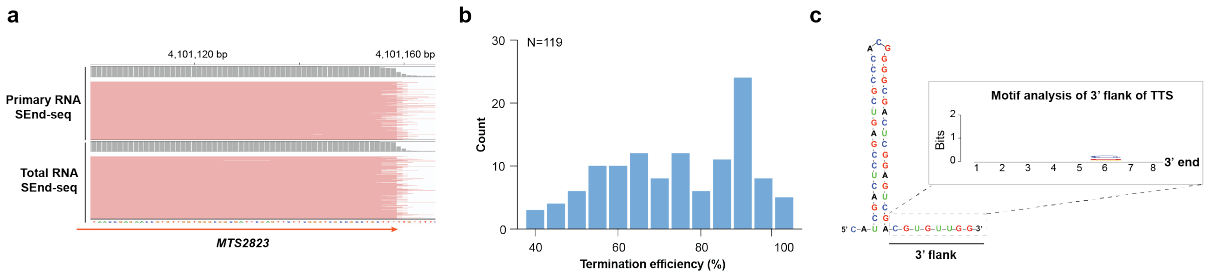


**Extended Data Figure 3. Characterization of Mtb TTSs detected by SEnd-seq. a**, SEnd-seq data track showing a TTS in Mtb. **b**, Histogram showing the distribution of termination efficiencies for the 119 identified TTSs in Mtb. The lower bound set to call a TTS was 40%. **c**, Secondary RNA structure for the TTS shown in **a** and motif analysis for the 3' flanking sequences of all identified TTSs showing a lack of any conserved motif.

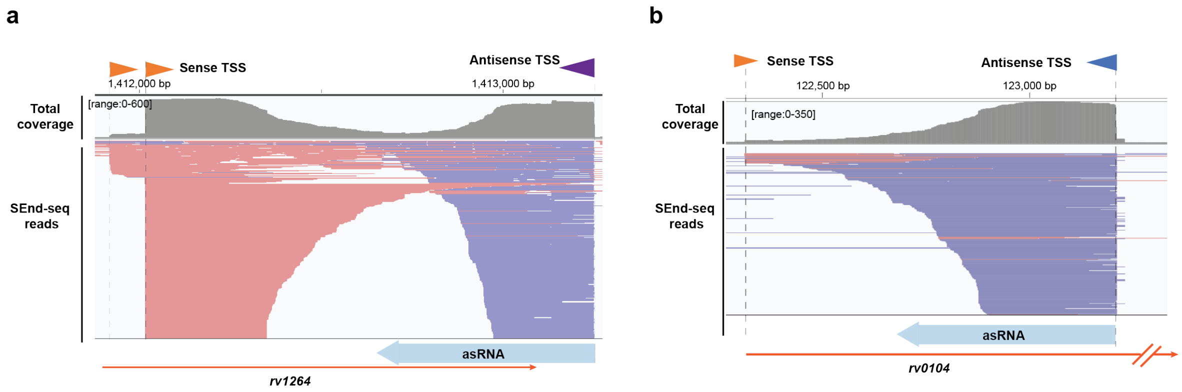


**Extended Data Figure 4. Pervasive antisense transcripts in Mtb detected by SEnd-seq. a-b**, SEnd-seq data tracks for two example Mtb genomic regions showing the abundance of antisense RNAs (blue lines).

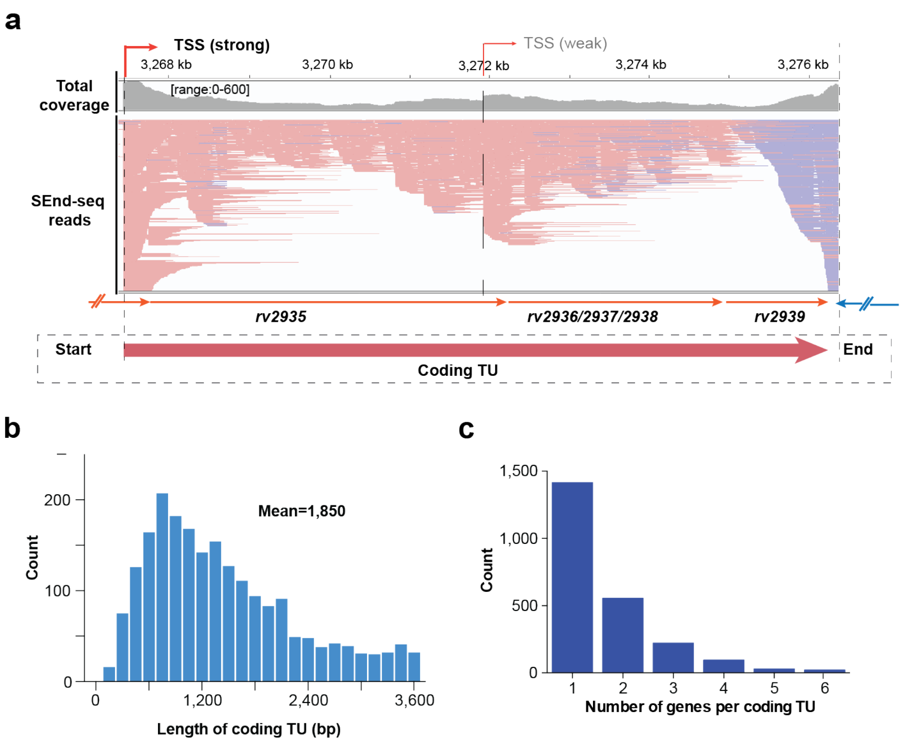


**Extended Data Figure 5. Characterization of coding transcription units (TUs) in Mtb. a**, SEnd-seq data track for an example TU consisting of multiple co-directional genes. This TU contains an upstream strong TSS and a downstream weak TSS. Thus, it was not further segmented. **b**, Distribution of the length of Mtb coding TUs. **c**, Distribution of the number of annotated genes within each coding TU.

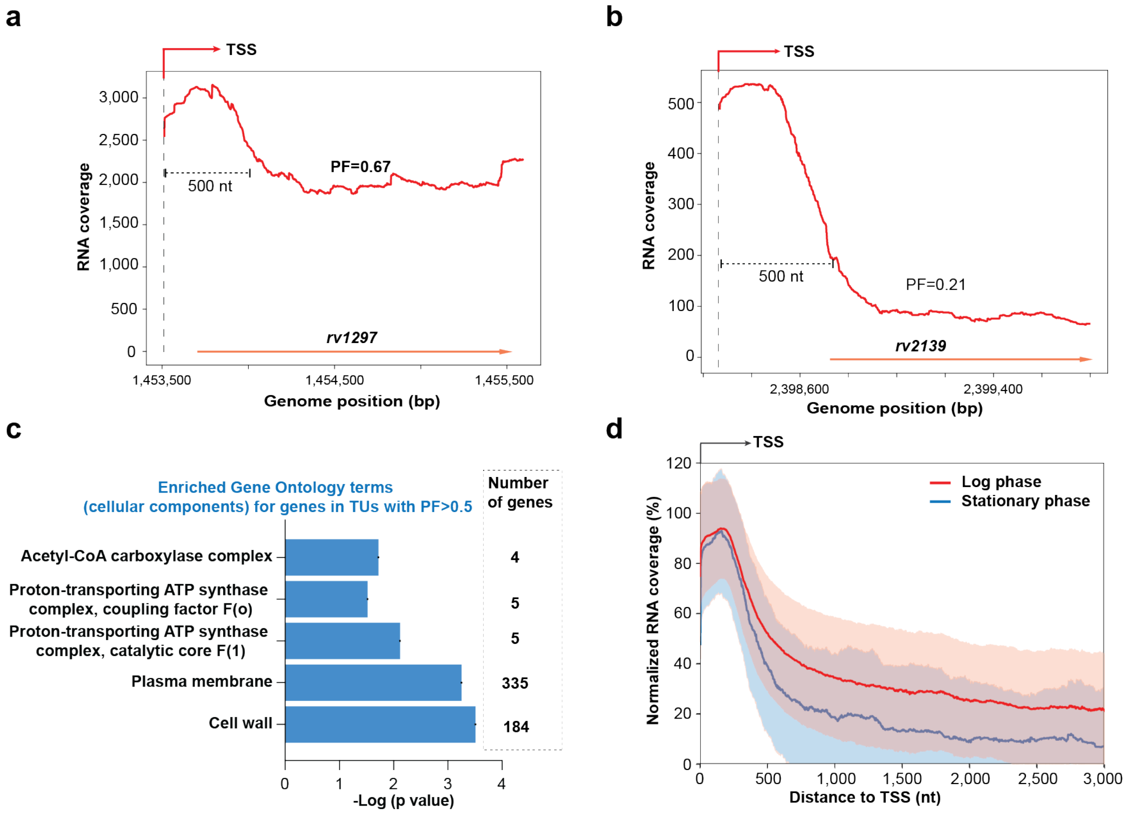


**Extended Data Figure 6. PF analysis of Mtb coding TUs. a**, SEnd-seq signals for an example TU (encoding the *rho* gene) with a relatively high PF. **b**, SEnd-seq signals for an example TU with a relatively low PF. **c**, Gene Ontology analysis of Mtb genes in TUs with a relatively high PF (>0.5). **d**, Summed RNA coverage aligned at TSSs for log-phase (red; 1,494 sites) and stationary-phase (blue; 302 sites) Mtb cells. RNA coverage was normalized to the peak value of each TU. Colored lines represent median values and shaded regions represent standard deviations.

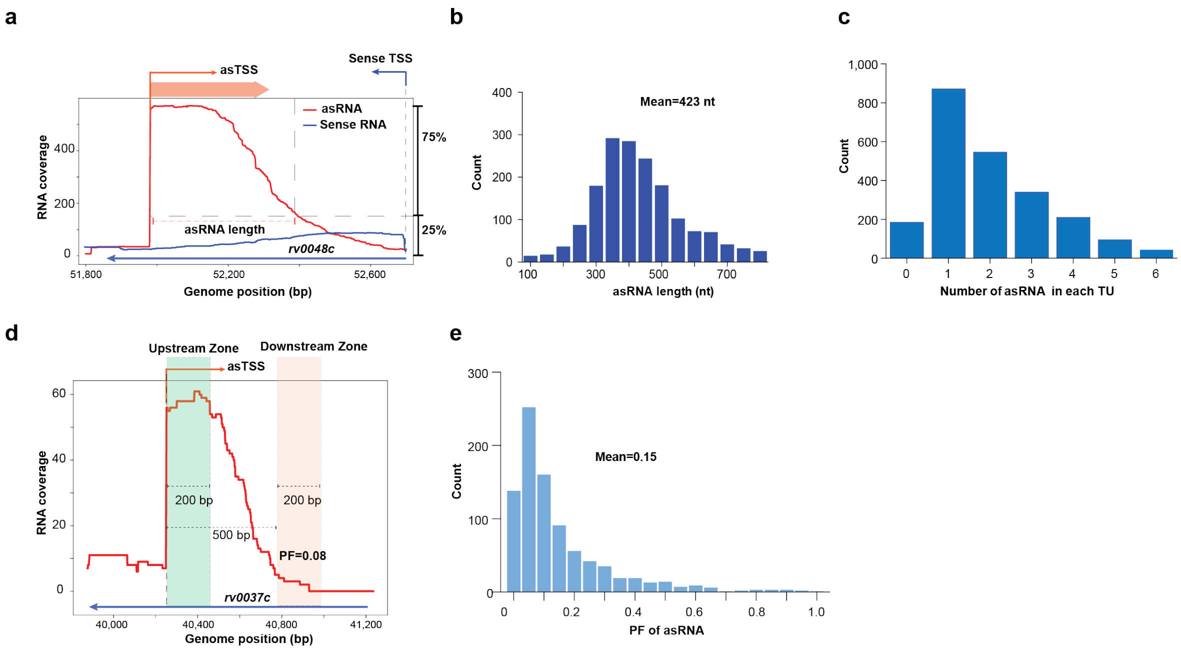


**Extended Data Figure 7. Characterization of asRNAs in the Mtb transcriptome. a**, SEnd-seq signals for an example asRNA and its corresponding sense RNA. **b**, Distribution of the length of asRNAs in log-phase Mtb cells. **c**, Distribution of the number of asRNAs within each coding TU. **d**, SEnd-seq signals for an example asRNA demonstrating the definition of upstream zone and downstream zone for PF calculation. **e**, Distribution of PF values for asRNAs from log-phase Mtb cells.

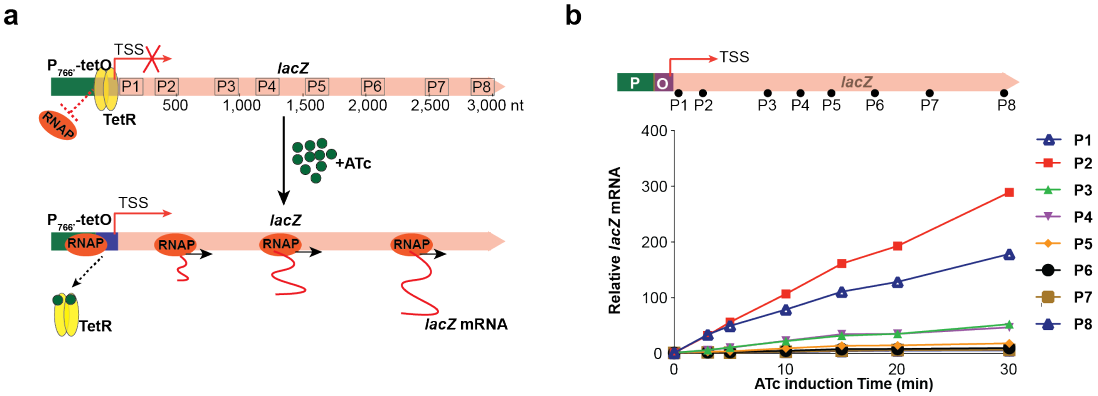


**Extended Data Figure 8. Measurement of heterologous *lacZ* transcription in Mtb. a**, Experimental scheme for measuring the transcriptional output of chromosome-integrated *lacZ* in Mtb cells induced by ATc. **b**, qPCR measurements on the kinetics of mRNA accumulation upon ATc induction at different locations (P1 to P8) across the *lacZ* gene body.

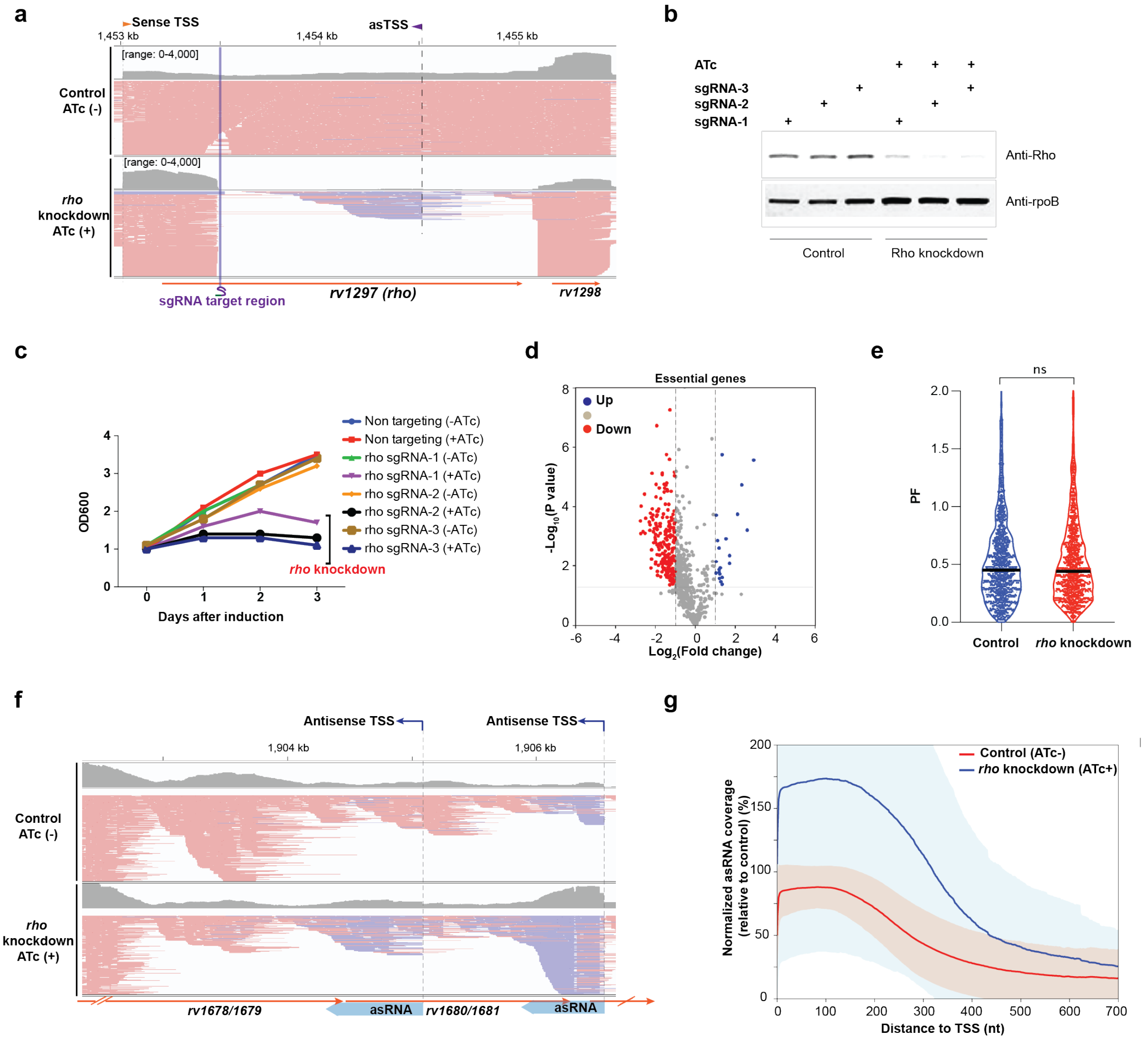


**Extended Data Figure 9. Effect of *rho* knockdown on Mtb gene expression.** **a**, SEnd-seq data track for Mtb cells with or without ATc-induced sgRNA expression showing the successful knockdown of *rho*. The sgRNA targeting site is indicated. **b**, Depletion of Rho proteins upon *rho* knockdown verified by western blotting. The RpoB signal serves as a loading control. **c**, Mtb growth curves with or without ATc-induced *rho* knockdown. **d**, Volcano plot showing changes in the expression of 696 essential Mtb genes upon *rho* knockdown measured by SEnd-seq. **e**, Violin plot comparing the distribution of PFs for coding TUs with or without *rho* knockdown. **f**, SEnd-seq data track for an example Mtb genomic region showing the increase in asRNA expression upon *rho* knockdown. **g**, Summed coverage for Mtb asRNAs aligned at TSSs showing the increase in asRNA abundance upon *rho* knockdown. RNA coverage of each asRNA was normalized to the peak value in the control group. Colored lines represent median values and shaded regions represent standard deviations.

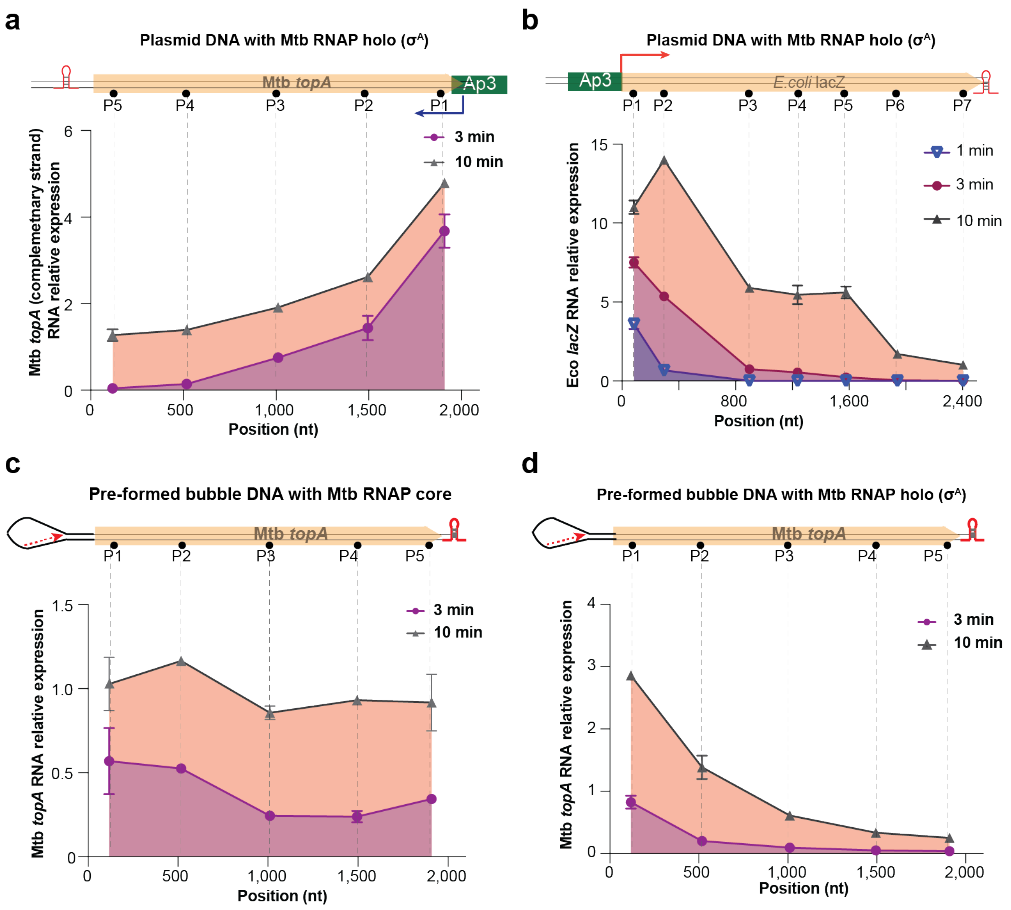


**Extended Data Figure 10. Characterization of the RNA products from in vitro transcription assays.** **a**, Length profile of RNA products measured by qPCR using Mtb RNAP/$\sigma^A$ holoenzyme and the same DNA template as in **Fig. 3c** but initiated from the opposite strand. **b**, Length profile of RNA products using Mtb RNAP/$\sigma^A$ holoenzyme and a different plasmid DNA template containing the Eco *lacZ* gene body. **c**, Length profile of RNA products using Mtb RNAP core enzyme (without $\sigma^A$) and a DNA template containing a pre-formed transcription bubble and an RNA primer. **d**, Same as **c** except that Mtb RNAP/$\sigma^A$ holoenzyme was used.

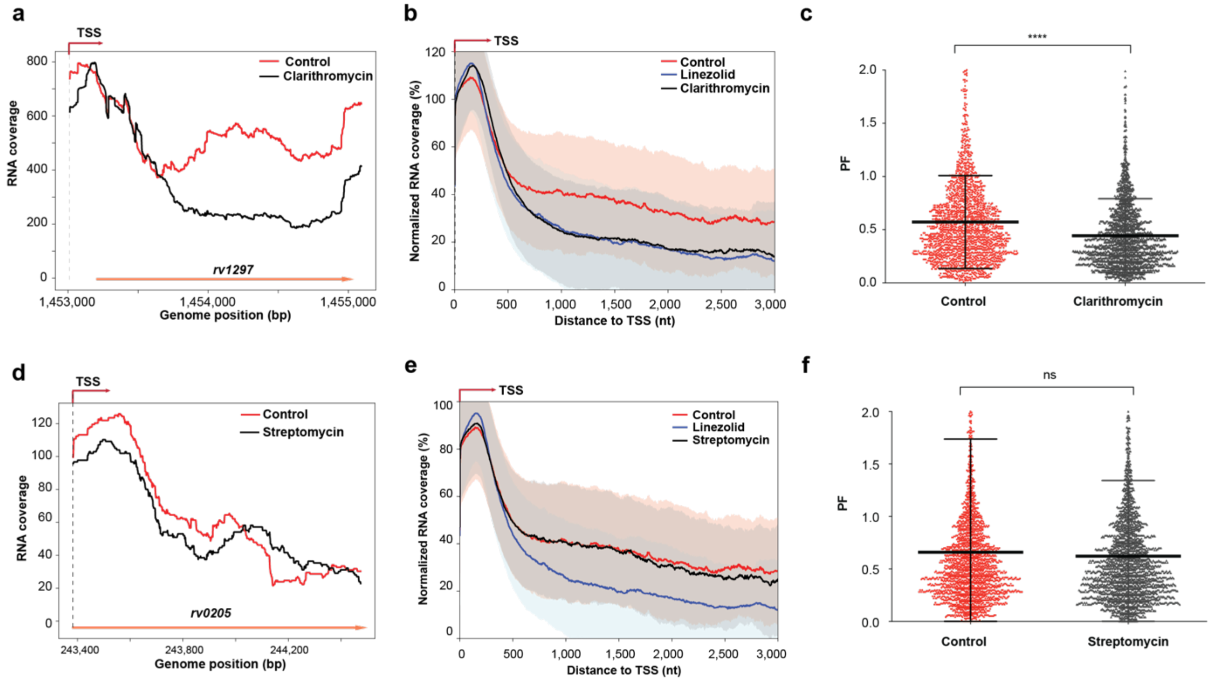


**Extended Data Figure 11. Effects of translation inhibitors on the processivity of Mtb transcription. a**, SEnd-seq signals in an example Mtb TU showing the difference in RNA coverage between the clarithromycin-treated condition (black) and the DMSO-treated control condition (red). **b**, Summed RNA coverage in TUs aligned at TSSs for DMSO-treated (control; red), linezolid-treated (blue), and clarithromycin-treated (black) Mtb cells. RNA coverage of each TU was normalized to the peak value in the control group. Colored lines represent median values and shaded regions represent standard deviations. **c**, Distribution of PFs for TUs in Mtb cells treated with DMSO (control) or clarithromycin. The black bars indicate mean values and standard deviations. **d**, SEnd-seq signals in an example Mtb TU comparing the RNA coverage between the streptomycin-treated condition (black) and the DMSO-treated control condition (red). **e**, Summed RNA coverage in TUs aligned at TSSs for DMSO-treated (control; red), linezolid-treated (blue), and streptomycin-treated (black) Mtb cells. **f**, Distribution of PFs for TUs in Mtb cells treated with DMSO (control) or streptomycin. The black bars indicate mean values and standard deviations.

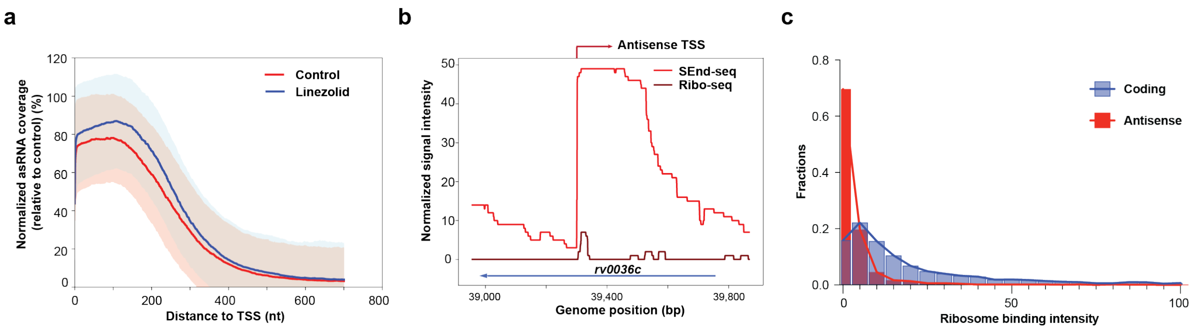


**Extended Data Figure 12. Role of translation in Mtb antisense transcription. a**, Summed coverage for Mtb asRNAs aligned at TSSs from DMSO-treated (control; red) and linezolid-treated (blue) cells. Colored lines represent median values and shaded regions represent standard deviations. **b**, SEnd-seq and Ribo-seq signals (reporting RNA coverage and ribosome binding, respectively) in an Mtb genomic region containing an antisense TSS. **c**, Distribution of ribosome binding intensities within Mtb coding TUs and asRNA regions (normalized by their lengths).
